# Spatial and ontogenetic variation in susceptibility to polarotacticecological traps

**DOI:** 10.1101/282046

**Authors:** Giovanna Villalobos-Jiménez, Rochelle Meah, Christopher Hassall

**Author notes:** Corresponding author. ORCiD: 0000-0003-3248-3644. These authors contributed equally to the work (Giovanna Villalobos-Jiménez, Rochelle Meah).

## Abstract

Ecological traps occur when environmental cues become unreliable, causing an evolutionary mismatch between features of the environment and expected outcome that leads to suboptimal behavioural responses and, ultimately, reduced fitness. Ecological traps arise due to anthropogenic disturbance in the environment introducing novel elements that mimic those environmental cues. Therefore, ecological traps represent a strong selective pressure in areas where anthropogenic changes are frequent, such as cities. However, given the exposure to these traps over generations, localised adaptations to ecological traps might be expected in urban populations. Dragonflies and damselflies (Odonata) are one of the many taxa vulnerable to ecological traps: odonates use horizontally polarised light as a cue of suitable water bodies, although some artificial surfaces also reflect horizontally polarised light strongly, thus misleading odonates to oviposit preferentially on these unsuitable surfaces rather than in water. Here, we compare the behavioural response to horizontally polarised light between urban and rural populations of the odonate *Ischnura elegans* to test the potential for localised adaptations to ecological traps. Laboratory choice experiments were performed using field-caught adults from urban and rural areas, and individuals reared in controlled conditions to account for environmental variation and exposure to polarised light. We also studied the association between ontogeny and polarotaxis that has been suggested – but not empirically tested – by other studies. The results showed that field-caught rural individuals had a significantly stronger preference for horizontally polarised light compared to urban individuals, suggesting there is strong selection against polarotaxis in urban areas. However, individuals reared in controlled conditions showed no difference between urban and rural populations, suggesting that there has not yet been adaptation in urban odonates. Instead, adults developed a strong preference for horizontally polarised light with increasing age, showing that mature adults are more prone to ecological traps. Possible mechanisms driving this response are discussed.

## Introduction

Organisms have evolved to recognise cues in the environment that guide their behaviour to maximise fitness (Orians and Wittenberger 1991). Decision-making based on these cues remains adaptive when, over evolutionary time, these cues exhibit stable congruence with survival and reproductive success (Hutto 1985). Ecological traps occur when environmental cues used by organisms for habitat choice are no longer reliable, such as when rapid environmental change decouples a behavioural cue from a fitness benefit, leading to maladaptive behaviour (Schlaepfer et al. 2002). Land use change, such as during agricultural expansion or urbanisation, is the most significant anthropogenic impact on terrestrial biodiversity (Sala et al. 2000) and is continuing at a rapid pace (Goldewijk 2001), making ecological traps increasingly frequent. The term was first coined by Gates and Gyles (1978) to describe the preference of passerines for nesting in narrow field-forest edges induced by anthropogenic disturbance, despite the fact that eggs and nestlings were more susceptible to predation and nest parasitism due to density-dependent mortality. However, the first experimental evidence to support the fact that artificial forest edges act as ecological traps was provided by Weldon and Haddad (2005). Using a landscape-scale experiment composed of habitat patches of equal area that differed in their amount of edge, they demonstrated that the amount of edge can adversely affect nest site selection and reproductive success of *Passerina cyanea*. The reason why *P. cyanea* prefers artificial forest edges despite the reduction in fitness becomes evident once their criteria for habitat selection are analysed. Forest edges are heterogeneous and provide a wide variety of resources for mixed-habitat birds (Gates and Gysel 1978, Weldon and Haddad 2005). On the other hand, artificial field-forest edges are much narrower and yet show the same heterogeneity and variety of structural cues of these species for settling and nesting (Weldon and Haddad 2005), and therefore are perceived as a suitable habitat. However, narrow forest edges increase bird density by concentrating nests, which in turn increases density-dependent mortality through predation and parasitism (Weldon and Haddad 2005). From an evolutionary perspective, anthropogenic field-forest edges are recent, thus passerines have not evolved to associate narrow man-made forest edges with increased predation and parasitism (Gates and Gysel 1978), if they are able to do so at all. The species becomes constrained by its evolutionary past to choose poorly, despite having suitable conditions available elsewhere, leading to reduced fitness (Schlaepfer et al. 2002).

Another group of organisms known to suffer from ecological traps are semi-aquatic insects. These organisms are attracted to horizontally polarised light since it is one of the main cues for detecting suitable water bodies (Bernáth et al. 2002). However, many surfaces also reflect horizontally polarised light, such as crude oil ponds (Horváth and Zeil 1996, Horváth et al. 1998), car bodies (Wildermuth and Horváth 2005, Kriska et al. 2006, Blahó et al. 2014), asphalt (Kriska et al. 1998), gravestones (Horváth et al. 2007), and solar panels (Horváth et al. 2010). In addition, these surfaces can reflect horizontally polarised light more strongly than water, making them potentially more attractive (Horváth et al. 1998). This leads to insects drowning in oil pools or ovipositing on other unsuitable surfaces instead of ovipositing on water, thus drying out the eggs and ultimately reducing fitness.

Polarised light pollution (PLP) has been identified as a novel source of environmental pollution (Horváth et al. 2009), but there is a significant lack of understanding concerning the strength, mechanism, and evolution of polarotaxis that is needed to mitigate the negative effects of polarotactic ecological traps (Robertson et al. 2013). Studies instead focus on measuring traps in the field with inferences about selective pressures (Robertson and Hutto 2006, Robertson et al. 2013). These studies contribute to understanding the impact of ecological traps, but they do not account for behavioural plasticity, therefore it is difficult to ascertain whether the (now) maladaptive behaviour is somehow dictated genetically – and thus passed over generations – or if it can be decoupled via plasticity.

The deleterious impacts of PLP, causing animals to misjudge suitable habitats for reproduction and dispersal, can result in significant impacts on populations (Horváth et al. 2009). PLP represents a strong selective pressure particularly upon urban populations considering many artificial surfaces that reflect horizontally polarised light are common in cities (Horváth et al. 2009, Faeth et al. 2012, Villalobos-Jiménez et al. 2016). If a given population does not have the behavioural plasticity to adapt to novel environmental cues, or if the population density is too small to persist until adaptation can occur, ecological traps can lead to extensive maladaptive behaviours resulting in population declines and extirpations (Schlaepfer et al. 2002).

A further complexity to the response of odonates to PLP is that preference to horizontally polarised light may vary according to ontogeny, since immature odonates tend to disperse away from water bodies until they reach maturity, and then return to breed (Wildermuth 1998, Horváth et al. 2007). Sexually immature adults are thought to exhibit negative polarotaxis (i.e. they actively avoid horizontally polarised light), but when these individuals reach sexual maturity, they become attracted to horizontally polarised light instead (Corbet 1999). Thus, sexual maturation may modulate the behavioural response to horizontally polarised light in odonates. However, this ontogenetic shift in the response to horizontally polarised light has not been empirically tested, and if present, then susceptibility to PLP could vary amongst age groups in odonate populations. In addition, if some age groups show differences in polarotaxis, this variation could provide a potential route for species to adapt in the face of PLP-induced reductions in fitness.

Here, we test local adaptations to ecological traps in odonates associated with PLP. We examine experimentally the behavioural response of the odonate *Ischnura elegans* to horizontally polarised light across different ages in the adult phase and compare between urban and rural populations. Contrary to other studies, we use wild-caught adults to study their response in the field, as well as laboratory rearing to control for plasticity and examine the ontogeny of the trait. Given that odonates in urban areas are more prone to exposure to PLP over generations, we hypothesize that individuals from more urbanised populations will exhibit reduced preference or even avoidance to horizontally polarised light to avoid reduced fitness. We expect the same response from both field-caught and laboratory-reared adults, except if other forces such as natural selection are present. Additionally, we expect an ontogenetic shift in the response to horizontally polarised light, with sexually immature individuals avoiding horizontally polarised light, but sexually mature individuals showing a strong preference to polarised light.

## Materials and methods

### Collection of adults and larvae rearing

We used individuals from common rearing conditions to evaluate the ontogenetic variation in the polarotactic response in adult odonates. We obtained *I. elegans* eggs from field-caught adult females from 3 urban and 5 rural sites (Table 1) during July-August 2015. Classification of sampling sites was according to the proportion of urbanisation within 1 Km around each pond, which was calculated using the 25m Land Cover Map 2007 (Natural Environment Research Council, Centre for Ecology & Hydrology, www.ceh.ac.uk/services/land-cover-map-2007) in ArcGIS 10.1 (ESRI 2011). Sites with an urban cover of >45% in a 1 Km buffer were classified as urban, whereas sites with an urban and suburban cover of <45% in a 1 Km buffer were classified as rural. Once the eggs hatched, the larvae were reared at 20°C at a photoperiod of 14L:10D with aerated tap water and fed with *Artemia* sp. and *Daphnia magna ad libitum*. Once the larvae emerged, the adults were kept in an insect net cage and fed with *Drosophila melanogaster ad libitum*. However, due to a large mortality rate in the larvae and low numbers of emerged adults reaching sexual maturity in the lab – particularly males – some of the individuals tested were collected as larvae from the field instead of being reared in the laboratory since the egg stage (reared from the egg stage: *N*_*urban*_ = 9, *N*_*rural*_ = 28; reared from the larval stage: *N*_*urban*_ = 36, *N*_*rural*_ = 10). Therefore, the individuals used in the tests that were reared in the laboratory are part of a semi-controlled rearing experiment (*N*_*urban*_ = 45, *N*_*rural*_ = 38).

**Table 1.**
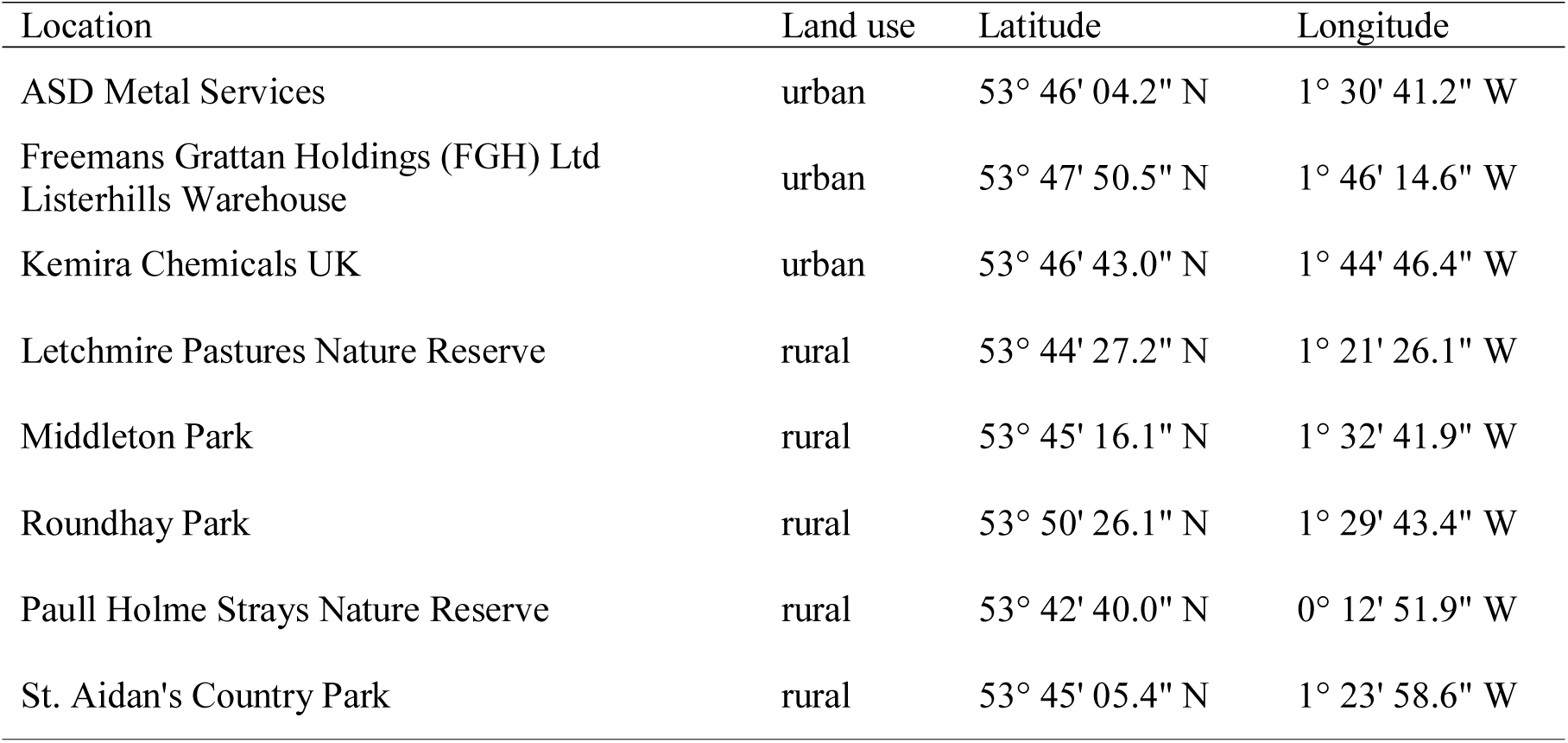
Sampling sites used for collecting *Ischnura elegans*. Land use classification of sampling sites was based on the percentage of urban cover (according to the Land Cover Map (LCM) 2007) at a distance of 1 Km around the ponds.

| Location | Land use | Latitude | Longitude |
| --- | --- | --- | --- |
| ASD Metal Services | urban | 53° 46' 04.2" N | 1° 30' 41.2" W |
| Freemans Grattan Holdings (FGH) Ltd<br>Listerhills Warehouse | urban | 53° 47' 50.5" N | 1° 46' 14.6" W |
| Kemira Chemicals UK | urban | 53° 46' 43.0" N | 1° 44' 46.4" W |
| Letchmire Pastures Nature Reserve | rural | 53° 44' 27.2" N | 1° 21' 26.1" W |
| Middleton Park | rural | 53° 45' 16.1" N | 1° 32' 41.9" W |
| Roundhay Park | rural | 53° 50' 26.1" N | 1° 29' 43.4" W |
| Paull Holme Strays Nature Reserve | rural | 53° 42' 40.0" N | 0° 12' 51.9" W |
| St. Aidan's Country Park | rural | 53° 45' 05.4" N | 1° 23' 58.6" W |

We also captured adult *I. elegans* from the field to compare the results with the lab-reared individuals (*N*_*urban*_ = 21, *N*_*rural*_ = 24). In this case, it was not possible to control for the mating status of the individuals (virgin *vs*. mated) or to analyse the changes in different ages since we did not know the exact date these odonates emerged, but instead we tested the response of only sexually mature individuals to polarised light. One of the advantages of using *I. elegans* is that sexually mature adults – both males and females – have a different colouration to their immature counterparts, thus the identification of mature adults in the field was based on their colouration. Also, immature *I. elegans* are rarely seen at the mating sites where samples were collected. The field-caught adult *Ischnura elegans* were placed in a resealable plastic bag within a cooler box to avoid heat stress while being transferred to the laboratory. On arrival, individuals were kept in an insect cage at 20°C and were fed with *D. melanogaster ad libitum*.

### Laboratory choice experiments

For the experiments, the adults were introduced into a dual-choice chamber made of matte cardboard (57 cm x 30 cm; Fig. 1). To control for the degree of polarisation, each individual was tested three times under three different conditions: (1) horizontally polarised light vs. unpolarised light (H-U test); (2) horizontally polarised light vs. vertically polarised light (H-V test); and (3) vertically polarised light vs. unpolarised light (V-U test). However, throughout this study we will only focus on the H-U test as this comparison provides the clearest test of the difference in response between the horizontally-polarised light characteristic of PLP and unpolarised light (to which odonates are also attracted). The results of the H-V and V-U tests are presented in the Supplementary Information. Polarised light was created using a linearly polarising filter (DyanSun 58mm slim linear polarising filter) positioned to reflect the required angle of polarisation. We also conducted a pilot field multiple-choice experiment using four test surfaces of varying polarisation cues to test for localised adaptations to PLP in the field (see Supplementary Information for further details).

**Fig. 1.**
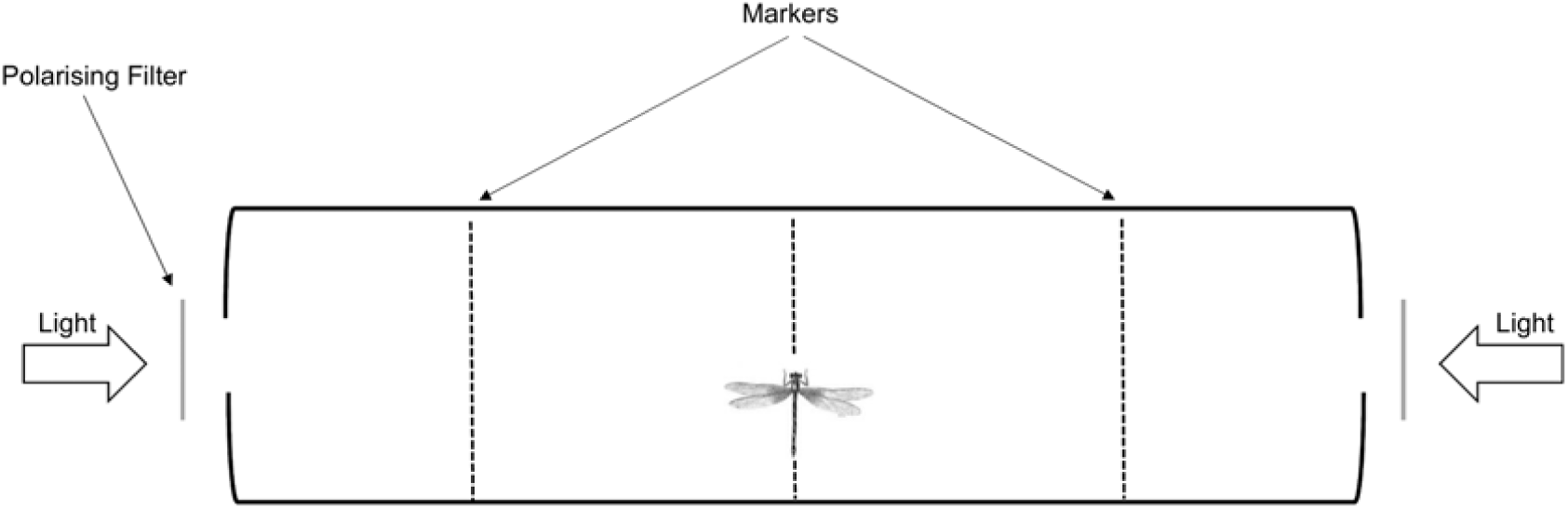
Experimental apparatus for testing individual preferences for polarised light.

To ensure a constant light intensity throughout the laboratory experiments, unfiltered light sources were covered by a semi-transparent material and measured using a light meter to within 10 lx of the light intensity of the filtered light source. The experiment was conducted to exclude any influence of external light surrounding the apparatus. Two markers were drawn intermediately between the centre of the apparatus and the two terminal light sources, at a distance of 14.5 cm from each light source (Fig. 1). Individuals were deemed to have chosen a preferred light source upon reaching either of the markers. The time taken for the individual to reach the marker was recorded as well as the position of the individual within the apparatus at 20, 40, and 60 seconds and then at 30 second intervals for 10 minutes or until the animal reached a marker. If no marker was reached within the 10 minutes of observation then the individual was deemed as showing no preference. To study the ontogeny of polarotaxis, the lab-reared individuals were tested every 2 days, starting from day 2 after emergence until they died. This allowed the evaluation of behavioural shifts across the adult phase. The field-caught individuals were only tested once since wild *I. elegans* cannot be aged reliably in the field, so we consider the response in this experiment to be representative only of sexually mature individuals.

### Statistical analyses

All statistical analyses of laboratory preference experiments were performed using R 3.0.1 (R Core Team 2013). To test the differences in the behavioural response to polarised light across different ages in urban and rural populations, we used a Generalised Linear Mixed Model (GLMM) with binomial error using the lme4 package (Bates et al. 2014), where light choice in each type of test used in the experimental setup was the response variable, age and land use (urban and rural) were used as explanatory variables and individual ID was the random effect to account for repeated measurements. However, in field-caught specimens, we used only a Generalised Linear Model (GLM) with binomial distribution, since all measurements were independent. In this model, the light choice was also the response variable, but only land use was the explanatory variable. Initially, sex was also considered as an explanatory variable for the analyses with individuals reared in the laboratory as well as the field-caught adults. However, sex did not contribute significantly to the models and was therefore eliminated.

We also tested the latency of choice making as a measurement of the strength of attraction to polarised light across different ages in urban and rural populations. In this case, we used the Cox proportional hazards method using the survival package (Therneau 2015) in R, with time of decision-making and light choice as the response variables, land use and age as explanatory variables, and clustering the data by the individual ID to account for repeated measurements in lab-reared odonates. With the field-caught specimens, only land use was the explanatory variable and clustering of data was not necessary since the measurements here are independent. Once again, sex was not included as an explanatory variable in these analyses given that it did not contribute significantly to the models.

## Results

When the preference for horizontally polarised light against unpolarised light was analysed in wild-caught adults, individuals caught from urban areas had a significantly lower probability of selecting horizontally polarised light compared to rural individuals (β = -2.023, SE = 0.898, *z* = -2.252, *P =* 0.024; Fig. 2a). Regarding the latency of decision-making, urban individuals took significantly longer to choose horizontally polarised light compared to rural individuals (β = -1.303, Hazard Rate (HR) = 0.272, SE = 0.656, *z =* -1.984, *P* = 0.047; Fig. 3a). Results of H-V test and V-U test are in Supplementary Information.

**Fig. 2.**
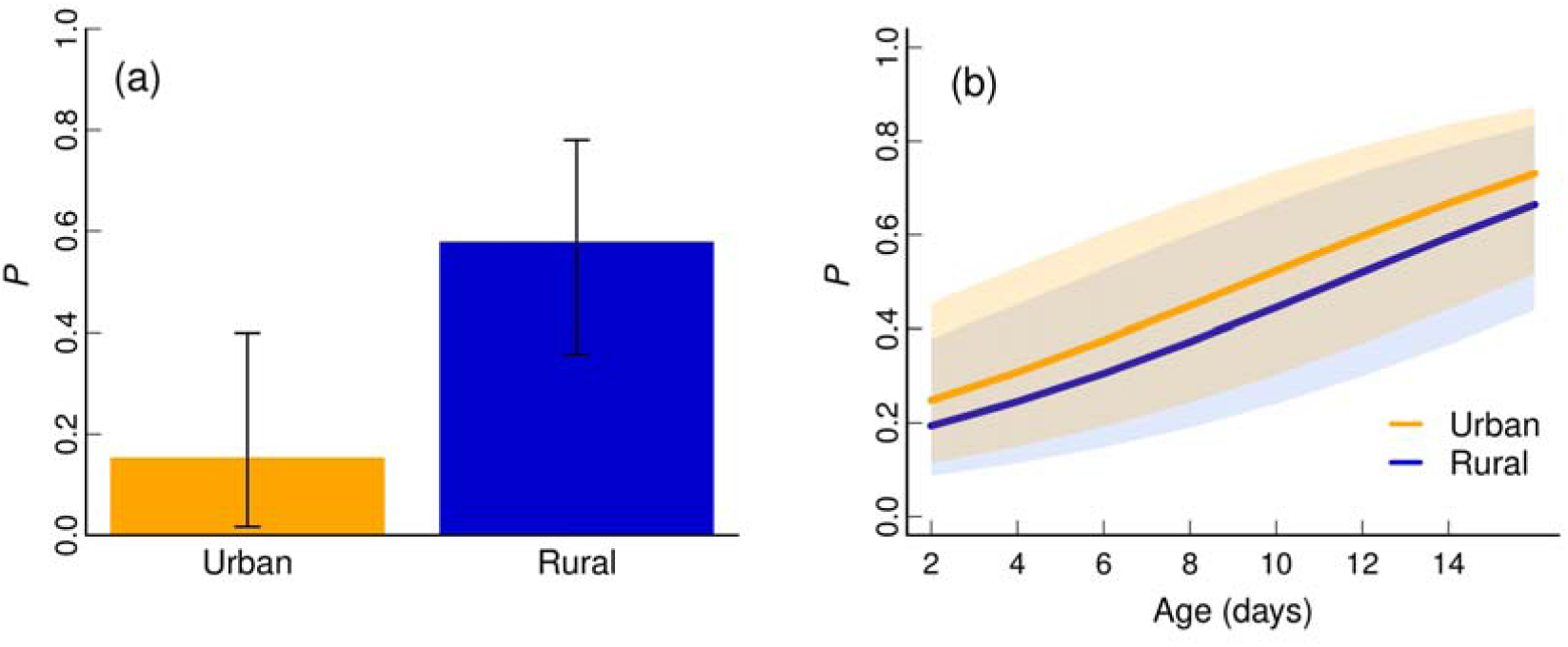
Preference for horizontally polarised light in H-U test between urban and rural populations: (a) the probability (*P*) of choosing horizontally polarised light in field-caught adults, error bars represent 95% CI; (b) The relationship between age and the probability (*P*) of choosing horizontally polarised light in adults from the common garden experiment. Shaded areas represent 95% CI.

**Fig. 3.**
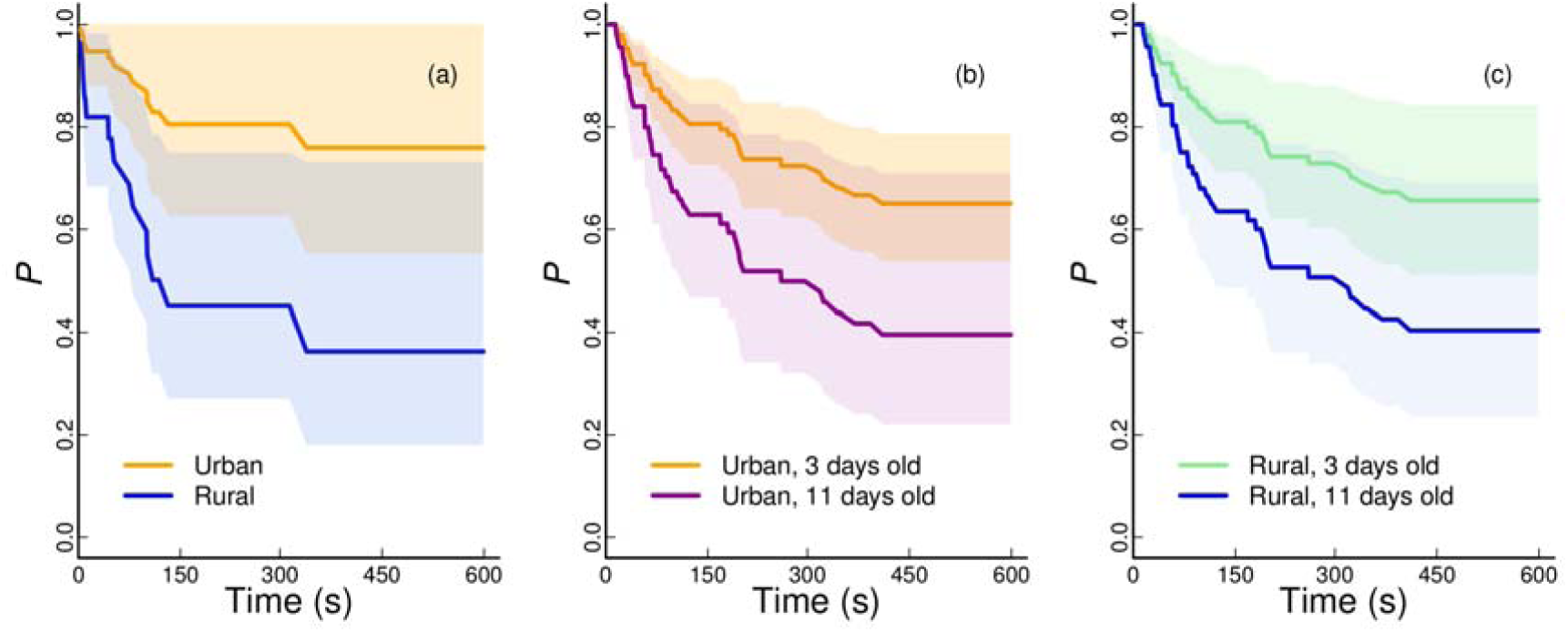
Latency of decision-making in H-U test between urban and rural individuals. showing the probability (*P*) of not choosing horizontally polarised light over time in (a) field-caught adults from urban and rural areas; (b) immature (3 days old) and mature (11 days old) urban individuals from the common rearing experiment; (c) immature (3 days old) and mature (11 days old) rural individuals from the common garden experiment. Shaded areas represent 95% CI.

On the other hand, the semi-controlled rearing experiment showed contrasting results to those obtained from wild-caught animals. Age exhibited a significant effect on the preference for horizontally polarised light (β = 0.151, SE = 0.068, *z* = 2.215, *P* = 0.027; Fig. 2b), with older individuals having a stronger preference for horizontally polarised light. However, the response of urban individuals reared in the laboratory was not significantly different to the response from rural individuals reared in the laboratory (β = 0.314, SE = 0.424, *z* = 0.741, *P* = 0.459; Fig. 2b), contrary to the results from the test using adults caught in the field. Regarding the latency of decision-making, age had a significant impact, with older individuals taking less time to choose horizontally polarised light (β = 0.096, HR = 1.100, SE = 0.043, *z* = 2.699, *P* = 0.008; Fig. 3b, c). On the other hand, no difference between urban and rural individuals was found in the time taken to choose (β = 0.022, HR = 1.022, SE = 0.326, *z =* 0.064, *P* = 0.949; Fig. 3). Results of H-V test and V-U test are in Supplementary Information.

## Discussion

Ecological traps represent a subtle and insidious threat to biodiversity, by reversing the relationship between environmental cues and fitness benefits. However, ours is the first study to explore the possibility of adaptation to ecological traps, as well as the ontogenetic shift in PLP risk in odonates. The results showed that field-caught individuals from urban areas have a significantly lower probability of selecting horizontally polarised light compared to individuals caught from rural areas. Likewise, field-caught individuals from urban areas took significantly longer to make a decision in this test. Unexpectedly, the results from the semi-controlled rearing experiment suggest that land use (urban or rural) does not influence the preference for horizontally polarised light in odonates when reared under common conditions. Instead, age has a significant impact on the behavioural response to horizontally polarised light, with older individuals preferring horizontally polarised light compared to non-polarised light. Age also had a significant impact in the latency of decision-making, with older individuals taking less time to choose horizontally polarised light compared to non-polarised light, though no significant difference between urban and rural individuals was found in this test. Below, we provide possible mechanisms driving the behavioural responses found to horizontally polarised light.

Preference for horizontally polarised light has developed in odonates as a cue for suitable habitat, which elicits reproductive behaviour such as oviposition in females or patrolling in males (Horváth and Varjú 1997, Wildermuth 1998, Bernáth et al. 2002). Our empirical demonstration of an ontogenetic switch from negative to positive polarotaxis fits with this theory that polarotaxis is associated with reproductive behaviour. An ontogenetic switch has also been found in other aquatic insects, such as mayflies (Szaz et al. 2015) and mud beetles (*Heterocerus* spp.) (Boda et al. 2014), and this switch has considerable implications for population-level responses to ecological traps since our results suggest mature individuals are more susceptible to ecological traps. There is also a consequence for our understanding of the mechanism underlying the response, since there may be physiological factors associated with sexual maturity that are potentially involved in polarotaxis, e.g. juvenile hormone (JH). JH is key in regulating a range of physiological, behavioural, and life-history traits such as mating, immune response, senescence, and survival (Córdoba-Aguilar and González-Tokman 2014). JH promotes egg production in females (Flatt et al. 2005) and increases territorial behaviour in males (Contreras-Garduño et al. 2009), but also promotes rapid senescence (González-Tokman et al. 2013) and decreases immune response (Contreras-Garduño et al. 2009). Thus, JH may play an important role in facilitating the polarotactic response of odonates, though the mechanism is unknown.

Importantly, we demonstrate a behavioural discrepancy between the individuals from the common rearing experiment and field-caught adults. The results from the lab-reared individuals suggest that there is no genetic basis underlying the decreased preference for horizontally polarised light found in odonates from urban areas, therefore there is no true adaptation. However, we suggest that the reduced response found in urban field-caught odonates arises from natural selection driving the removal by ecological traps of those animals that have a high natural polarotactic response. This hypothesis is supported by preliminary experiments conducted by supplying artificial PLP traps to odonates in the field, which suggested that rural traps collected more odonates regardless of local population densities (see Supplementary Information). Given that artificial horizontally-polarising cues are less frequent in rural areas (Horváth et al. 2009), there is no such bottleneck acting upon these populations, which may explain why rural field-caught adults showed a strong preference to polarised light. Hence, while there is no apparent adaptation, there is still strong selection already acting upon urban populations leading to the potential for future generations to adapt to ecological traps. However, ecological traps have existed for a relatively short time period which may be insufficient for adaptation to occur in a trait that has had such an important role in the semi-aquatic lifestyle of these insects (Gates and Gysel 1978, Schlaepfer et al. 2002). According to Schlaepfer *et al*. (2002), if populations cannot adapt rapidly to ecological traps or the population is too small to persist until adaptation occurs, ecological traps lead to abrupt population declines or even extinction. Therefore, urban odonate populations may act as sinks in wider metapopulations unless odonates can adapt promptly to ecological traps.

The results also showed that rural field-caught adults were significantly quicker in choosing horizontally polarised light compared to their urban counterparts. The aim of decision making is to evaluate the advantages and disadvantages of all the alternative options available in order to avoid or minimise risks (McFarland 1977). This action depends on many factors, both external (weather conditions, predators) and internal (prior experiences, weight, egg loads) (McFarland 1977). The time invested in decision-making is crucial for acquiring the best outcome possible with increased accuracy and minimal risks (Zenger and Fahle 1997, Skorupski et al. 2006). However, there may be a trade-off between latency of decision-making and the benefit of such decision (Skorupski et al. 2006, Chittka et al. 2009). For example, damselflies that take longer to decide to approach a potential mate may have increased accuracy and avoid risks such as predation, but on the other hand they lose more opportunities for mating (Hilfert-Rüppell 1999). In various circumstances, natural selection may be in favour of bold individuals, which take less time to make a decision despite the presence of risks (Kurvers et al. 2011). Many species that have been successful colonisers of urban areas exhibit increased boldness compared to other species (Evans et al. 2010, Lowry et al. 2013). However, in this case, individuals that take longer to make a decision are less susceptible to ecological traps, particularly in urban areas. There may be a trade-off between boldness in urban odonates and susceptibility to ecological traps. Thus, selective pressures may be already driving resistance to ecological traps found in field-caught urban odonates through enhanced latency to decision-making. However, as previously mentioned, these results were not found in lab-reared individuals, so there appears to be no adaptation yet.

On the other hand, individuals reared in the laboratory showed decreased latency of decision-making with increased age, which suggests there is a mechanism associated with sexual maturation and senescence that influences this behaviour. Studies have shown that testosterone in vertebrates, for instance, increases boldness (Raynaud and Schradin 2014) and also aggressiveness (Chang et al. 2012). The invertebrate homologue, JH, has shown similar results in damselflies (Contreras-Garduño et al. 2009). JH has been found to increase with age (González-Tokman et al. 2013), therefore it is within reason that older individuals took less time to make a decision. Even though it is very likely that JH is the main regulator driving the latency of decision-making in odonates (in polarotaxis and other behaviours), the empirical evidence supporting this idea does not exist to this date.

Despite the fact that urban and rural individuals reared in the lab showed no difference in their preference for horizontally-polarised light, there is indirect evidence which may suggest that urban odonate populations in the wild have a shorter life span compared to rural populations. The trailing edge of the flight period of odonates is shorter in cities than in rural areas, which suggests a contraction of the flight period and that the adult phase has a shorter life span (Villalobos-Jiménez and Hassall 2017). This may explain why rural individuals caught in the field showed a stronger preference for horizontally polarised light compared to the urban field-caught individuals.

To conclude, this study demonstrates that there are considerable differences in the susceptibility to polarised light pollution, and that these manifest more strongly in urban odonate populations. However, there is no evidence that this selection pressure has produced a genetic adaptation in urban populations. Ontogenetic variation in polarotaxis means that adult odonates are more susceptible to ecological traps than younger animals, which suggests that there may be subtle demographic effects on populations and implies a role for juvenile hormone in driving the response to polarised light. The findings from odonates may well extend to other semi-aquatic insects which exhibit similar behavioural responses to PLP, with significant consequences for the aquatic and terrestrial food webs in which such invertebrates play a fundamental role.

## Author contributions

The first and second author contributed equally to this paper.

Sample locations and results from the H-V and V-U tests are available in the Supplementary Information. Collected data and R analysis scripts are available in Figshare.

